# Highly inducible transcription is impaired in cells lacking Cyclin T1 (CycT1) of the positive transcription factor b (P-TEFb)

**DOI:** 10.64898/2026.09.26.754677

**Authors:** Takuya Kajitani, Fang Huang, Kenji Nishiura, Dan Irwin, Aung Chein, Koh Fujinaga

**Author notes:** Jiangxia Laboratory in Hubei, Wuhan 430000, China (Fang Huang).

## Abstract

In eukaryotic cells, the positive transcription elongation factor b (P-TEFb) plays a critical role in the transition of RNAPII from the paused state to actively transcribing mode. Although three different cyclin (Cyc) T (CycT1, T2a, and T2b) interact with CDK9 to form functional P-TEFb, a majority of P-TEFb complexes are comprised of the CycT1 and CDK9. To analyze the role of the CycT1 subunit of P-TEFb on cellular transcription, we established HEK293T cells lacking CycT1 proteins (CycT1-KO cells). While no apparent growth defects were observed with CYcT1-KO cells, the level of CDK9 was decreased and no compensatory over-expression of CycT2 was observed, suggesting that the reduced level of functional P-TEFb complexes is sufficient for the normal cell growth. Reporter gene assays indicate that NF-κB-dependent transcription induced by PMA was impaired in CycT1-KO cells although the function of NFκB (P65) *per se* was not affected. On the other hand, AP1-dependent transactivation induced by PMA was unaffected in CycT1-KO cells. Moreover, transcription stimulated by JQ1, a strong P-TEFb inducer, was reduced in CycT1-KO cells. Data of biochemical analysis indicate that P-TEFb with CycT1 (P-TEFb (CDK9:CycT1)) was efficiently released from 7SKsnRNP by JQ1 while P-TEFb (CDK9:CycT2) was not affected by JQ1. Transcriptome analysis indicated that the lack of CycT1 had a minor effect on the steady-state transcription. However, expression of JQ1-dependent genes was severely reduced in CycT1-KO cells. Interestingly, referring to gene expression profiles of 33 different types of cancer revealed that 11 out of 13 top JQ1-dependent genes were also highly upregulated in Pancreatic Adenocarcinoma. From these results, we conclude that although many genes can be regulated by both P-TEFb (CDK9:CycT1) and P-TEFb (CDK9:CycT2), CycT1 plays a critical role in regulating highly inducible genes, which are also aberrantly regulated in a particular type of cancer.

## Introduction

Eukaryotic transcriptional networks are highly orchestrated systems regulated by various factors including cell surface receptors, intracellular signalling molecules, nuclear transcription factors, epigenetic regulators, etc [1]. However, all regulatory pathways converge into a downstream on-off switch, represented by the pause-release of RNA polymerase II (RNAPII), that ultimately determines which genes are to be turned on or off [1]. On most protein-coding genes, RNAPII is engaged at the promoter region but paused immediately after the initiation of transcription [2]. RNAPII’s transition from a pausing to elongation is regulated by the positive transcription elongation factor b (P-TEFb) that phosphorylates the C-terminal domain (CTD) of RNAPII and negative transcription elongation factors [3–5].

P-TEFb is a heterodimer of cyclin-dependent kinase (CDK) 9 and one of three different cyclin(Cyc)s: CycT1, CycT2a, and CycT2b [6]. CycT1 and CycT2a/b are encoded by different genes (CCNT1 and CCNT2). CycT2a and CycT2b are produced by alternative splicing from the CCNT2 gene [6]. While the P-TEFb with CycT1 (P-TEFb (CDK9:CycT1)) seems to be a major form in many cell types, CycT2 plays an essential role during mouse embryogenesis, such as the development of skeletal muscles [7–9]. In cells, P-TEFb-dependent transcription is modulated by three different modes of regulation [3, 4]. Most studies on P-TEFb regulatory pathways have been done with P-TEFb (CycT1:CDK9), and whether P-TEFb (CycT2:CDK9) is regulated the same regulatory mechanisms has not been examined although it is assumed that CycT2 and CycT1 are inter-exchangeable because of the high sequence similarities between them.

First, protein levels of functional P-TEFb are strictly determined by the cellular status and the interaction between CDK9 and CycT1. While CDK9 proteins unbound to CycT1 are stably sequestered by the heat shock protein (HSP90), CycT1 proteins unbound to CDK9 are immediately degraded via ubiquitin-E3-ligase-dependent pathways [10, 11]. In resting T cells and unresponsive (anergic or exhausted) T cells, for example, CycT1 is dephosphorylated and dissociated from CDK9, resulting in the expression at very low level [12]. In normally replicating cells, CycT1 is phosphorylated by PKC and associated tightly with CDK9, resulting in high expression levels [12]. Second, even though functional P-TEFb proteins are abundantly expressed in normally replicating cells, large proportion (50-90%) of them are incorporated into 7SK small nuclear ribonucleoprotein (snRNP) complexes, where CDK9’s kinase activity is inhibited by 7SKsnRNA and HEXIM1/2 proteins [5]. Developmental and environmental stresses release P-TEFb from 7SKsnRNP and activate CDK9’s kinase activity [3, 13]. Third, “free” P-TEFb is also sequestered by Bromodomain Protein (Brd) 4 or recruited to the paused RNAPII via different recruitment factors including DNA-bound transcription factors, RNA-bound transcription factors, chromatin-associated factors, RNAPII-associated factors, the Super Elongation Complex, etc. All three modes of P-TEFb regulation are tightly linked with each other to fine tune the transcriptional networks [4].

Since many human diseases and conditions, including infectious diseases, immunosuppression, inflammation, auto-immune diseases, developmental disorders, and cancer, are associated with aberrant regulation of P-TEFb in cells, P-TEFb represents an excellent therapeutic target [13]. For example, currently many inhibitors for CDK9’s kinase activities are being tested in clinical trials as anti-cancer drugs [13]. However, since P-TEFb is required for transcriptional elongation of most genes, complete blocking of CDK9 will globally shut down cellular transcription and elicit severe side effects, which limits the usage of pan-CDK9 inhibitors with a very narrow therapeutic window. Therefore, it is necessary to control P-TEFb activities in a manner specific to its target genes.

While the mechanisms of individual P-TEFb-regulatory pathway have been extensively studied, little is known about how different sets of P-TEFb-target genes are transcribed upon different environmental or developmental stimuli. Moreover, it is largely unclear whether different P-TEFb complexes (CDK9:CycT1, CDK9:CycT2a, and CDK9:CycT2b) have distinct target specificities. To obtain information about the mechanisms by which P-TEFb’s target genes are determined, we genetically inactivated CycT1 to establish CycT1 knockout (KO) cell lines. Interestingly. CycT1-KO cells replicated normally compared with wild type (WT) cells, suggesting CycT2 can compensate the loss of CycT1 for normal cell replication. However, transcription activated by NFκB via PMA stimulation was impaired in CycT1-KO cells. Similarly, transcription stimulated by the direct induction of P-TEFb via an inhibition of the BET-Bromodomain with JQ1 was also reduced in CycT1-KO cells. Further analysis of P-TEFb induction by JQ1 revealed that P-TEFb (CDK9:CycT1) was efficiently released from 7SKsnRNP by JQ1 while P-TEFb (CDK9:CycT2) was not affected by JQ1. Global gene expression analysis indicates that the lack of CycT1 had a minor effect on the steady-state transcription while the expression of JQ1-dependent genes was severely reduced in CycT1-KO cells. Finally, searching gene expression profiles of 33 different types of cancer revealed that 11 out of 13 top JQ1-dependent genes were also highly upregulated in Pancreatic Adenocarcinoma. We conclude that CycT1 plays a critical role in regulating highly inducible genes, which are also aberrantly regulated in a particular type of cancer.

## Results

### Establishment and analysis of CycT1-KO cells

To analyze the role of the P-TEFb (CycT1:CDK9) on cellular transcription, CycT1 was genetically inactivated in HEK293T cells by CRISPR-Cas9. No differences in apparent cellular morphologies and rates of cell growth between wild-type 293T (WT) and CycT1 knockout (CycT1-KO) cells were observed (data not presented). Western blot analysis of CycT1-KO cells confirmed that no CycT1 proteins were detected (**Fig.1A**). No compensatory over-expression of CycT2 proteins was detected in these cells. However, the level of CDK9 proteins was reduced in CycT1-KO cells, suggesting that the amount of functional P-TEFb molecules was reduced concomitantly with the abolishment of CycT1 (**Fig.1A**). Co-immunoprecipitation with anti-CDK9 antibodies confirmed that CycT2 interacts with CDK9 to form P-TEFb complexes in CycT1-KO cells (**Fig.1B**, lanes 1 and 2). However, the interaction between CDK9 and HEXIM1 was reduced in CycT1-KO cells when the same amount of HEXIM1 was immunoprecipitated (**Fig.1B**, lanes 3 and 4), indicating that a larger portion of P-TEFb (CDK9:CycT2) complexes in these cells is not incorporated in the 7SKsnRNP. No differences in the protein levels of other 7SKsnRNP components including HEXIM1, LARP7, MEPCE were observed (**Fig.1A**), while a slight increase in the level of 7SKsnRNA in CycT1-KO cells were observed (**Fig.1C**). A similar attempt to establish CycT1 knockout cells in HeLa cells were unsuccessful, which might suggest that the lack of CycT1 is lethal in these cells.

**Figure 1.**
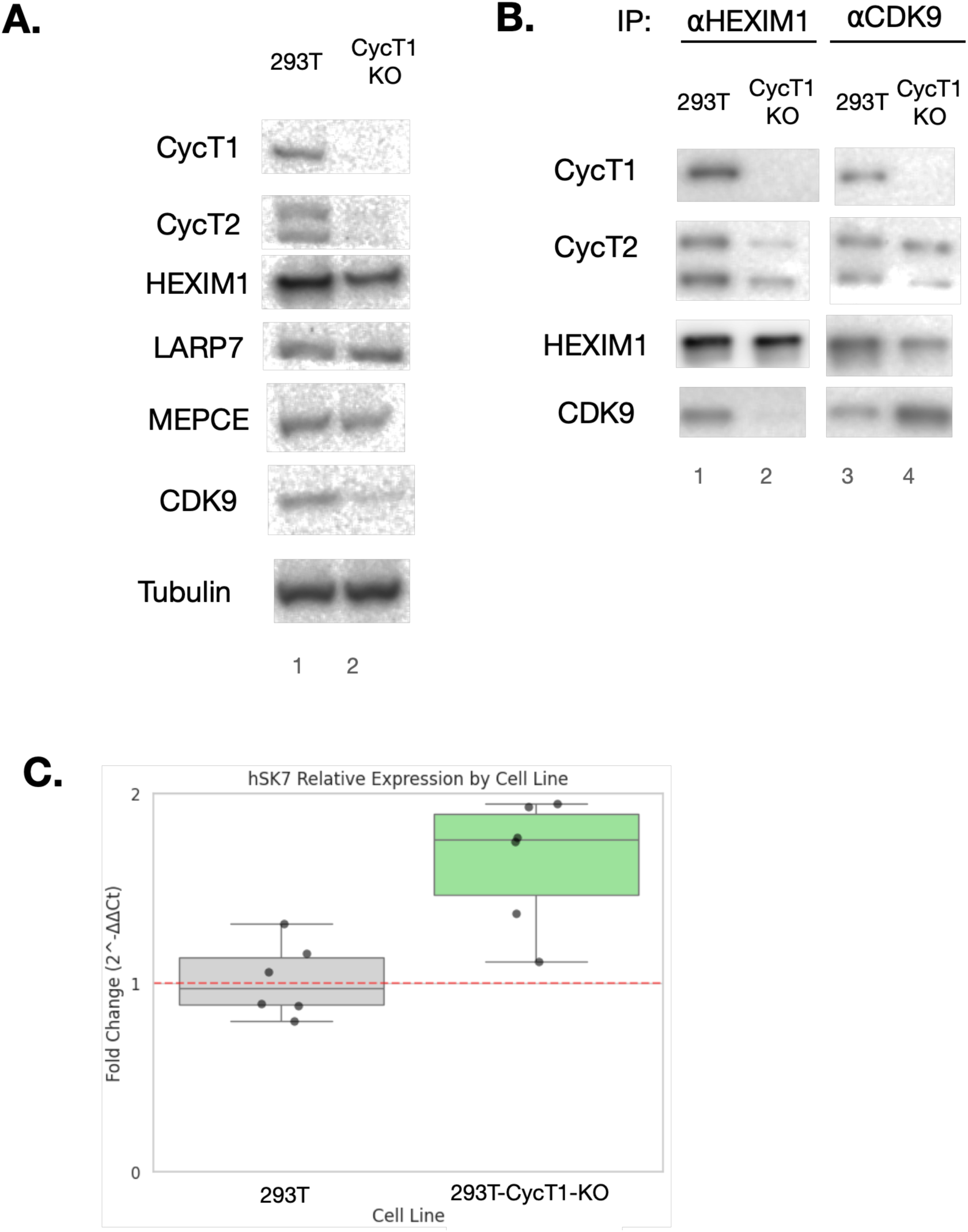
Characterization of CycT1-KO cells. **A.** 293T and CycT1-KO cells were lysed and the lysates were subjected to Western blotting using antibodies indicated at the left of each panel. *a*-Tubulin was used as a loading control. **B.** Lysates of 293T and CycT1-KO cells were subjected to co-immunoprecipitation with *a*-HEXIM1 (lanes 1 and 2) or *a*-CDK9 (lanes 3 and 4), followed by Western blotting using antibodies indicated at the left of each panel. Input samples are shown in **A. C.** Total cellular RNA was isolated from 293T and CycT1-KO cells and subjected to RT-qPCR analysis to quantify the expression levels of 7SKsnRNA. GAPDH mRNA levels were also measured and used to normalize the data.

### Transcription from inducible promoters is selectively impaired in CycT1-KO cells

Next, we examined whether the lack of CycT1 affects transcription by various well-defined reporter gene assays. First, since CycT1 is essential for the stimulation of transcription elongation by the Tat protein of the human immunodeficiency virus type 1 (HIV-1), WT and CycT1-KO cells were transfected with the reporter HIV-LTR-luciferase plasmids with or without co-expression of HIV-1 Tat proteins. While Tat stimulated the luciferase expression from HIV-LTR more than 10 folds in WT cells, it failed to stimulate the luciferase expression in CycT1-KO cells (**Fig.2A**, bars 1 and 2). Transient co-expression of CycT1 proteins stimulated Tat-dependent luciferase expression in CycT1-KO cells to the level similar to that in the WT cells (**Fig.2A**, bar 3). It was also demonstrated that the nuclear factor kappa B (NFκB) requires P-TEFb via interaction between the Rel A (P65) subunit and CycT1 [14]. Therefore, we next examined whether NFκB-dependent transcription is affected by the lack of CycT1. A reporter NFκB (P65)-luciferase plasmid was transfected, and cells were treated with PMA for 24 hours to induce NFκB. As shown in **Fig.2B**, PMA-dependent luciferase expression was significantly reduced in CycT1 KO cells. The ability of P65 to transactivate its target genes *per se* in CycT1-KO cells was examined by forced (artificial) tethering assays.

**Figure 2.**
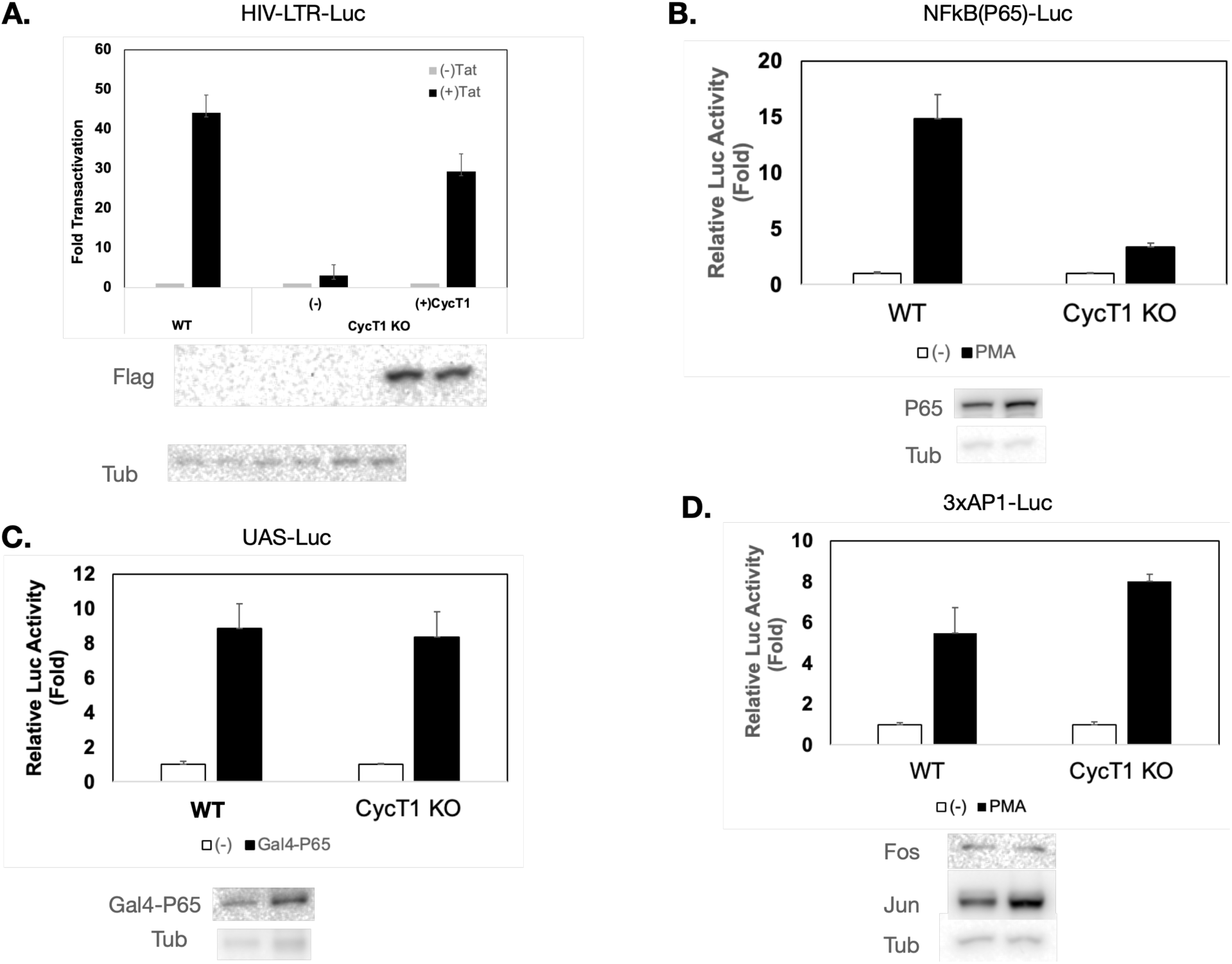
PMA-dependent NF-κB activity is impaired in CycT1-KO cells. **A.** 293T or CycT1-KO cells were co-transfected with HIV-LTR-luciferase reporter plasmid with or without Tat-expressing plasmid or CycT1-expressing plasmid. 48 hours after transfection, cells were harvested and subjected to luciferase assays. Results are presented as values relative to that obtained with 293T cells transfected with the reporter plasmid without ectopic expression of Tat and CycT1. (lower panels) Western blotting of the lysates confirmed the expression of ectopic expression. **B.** 293T or CycT1-KO cells were co-transfected with a luciferase reporter plasmid that containing NF-κB-binding sequences in the promoter region (NF-κB-Luc). 24 hours after transfection, cells were treated with DMSO or PMA for additional 24 hours. Cells were then harvested and subjected to luciferase assays. Results are presented as values relative to that obtained with 293T cells treated with DMSO. Western blotting confirmed that similar levels of NF-κB (P65) were expressed in these cells. **C.** 293T or CycT1-KO cells were co-transfected with a luciferase reporter plasmid that containing six Gal4-binding sequences in the promoter region (UAS-Luc) with or without the plasmid encoding the Gal4-P65 protein. 48 hours after transfection, cells were then harvested and subjected to luciferase assays. Results are presented as values relative to that obtained with 293T cells expressing no Gal4-P65. Western blotting confirmed that similar levels of Gal4-P65 proteins were expressed in these cells. **D.** 293T or CycT1-KO cells were co-transfected with a luciferase reporter plasmid that containing three AP-1-binding sequences in the promoter region (AP1-Luc). 24 hours after transfection, cells were treated with DMSO or PMA for additional 24 hours. Cells were then harvested and subjected to luciferase assays. Results are presented as values relative to that obtained with 293T cells treated with DMSO. Western blotting confirmed that similar levels of c-Jun and c-Fos proteins were expressed in these cells.

Gal4-P65 proteins activated UAS-Luciferase reporter genes in CycT1 KO cells to a level similar to that in WT cells (**Fig.2C**), suggesting that P65 can activate transcription when it is recruited to a promoter in CycT1-KO cells. Interestingly, the activity of another P-TEFb-dependent transactivator, AP-1, which can also be induced by PMA, was not affected in CycT1 KO cells (**Fig.2D**). These results indicate that CycT1 is required for selective transactivator-dependent transcription.

### Transcription stimulated by direct activation of P-TEFb is impaired in CycT1-KO cells

Next, we analyzed transcription activated by cellular stimulations that directly activate P-TEFb. We have previously demonstrated that BET-bromodomain inhibitors (BETi), such as JQ1, release P-TEFb from the 7SKsnRNP complex and stimulate P-TEFb activity [15]. In addition, we have also demonstrated that the *hexim1* gene is an immediate early gene responding to P-TEFb induction, and mapped the minimum *hexim 1* promoter that is activated by JQ1 (HexP) [16]. Therefore, we compared the effect of JQ1 on HexP-luciferase expression between WT cells and CycT1-KO cells. After transfection of the HexP-Luc reporter plasmid, cells were stimulated with JQ1 (5μM) for 24 hours prior to luciferase measurement. JQ1-dependent transactivation from HexP was significantly reduced (**Fig.2A**). Next, JQ1’s activity to release P-TEFb from 7SKsnRNP was examined by sequential salt extraction assays [17, 18]. P-TEFb in 7SKsnRNP is loosely associated with the nucleus and extracted with a low salt lysis buffer (LS: 10mM KCl) whereas released/activated P-TEFb is relocated into the chromatin fraction and extracted with a high salt lysis buffer (HS: 450mM KCl) [19]. In unstimulated WT cells, the level of CycT1 in the LS was similar to that in HS whereas the level of CycT2 in the LS was significantly lower than that in HS (**Fig.2B**, lanes 1 and 2), indicating that a larger portion of P-TEFb (CDK9:CycT2) is not incorporated in the 7SKsnRNP. Similarly in unstimulated CycT1-KO cells, the level of CycT2 in LS is significantly lower than that in HS (**Fig.2B**, lanes 5 and 6), which is consistent with the co-IP result (**Fig.1B**). The level of CycT1 in LS was reduced by JQ1 stimulation in WT cells (**Fig.2B**, lanes 1 and 3), suggesting that JQ1 releases P-TEFb (CDK9:CycT1) from 7SKsnRNP. However, the levels of CycT2 in both WT and CycT1-KO cells were only slightly reduced (Fig.2B, lanes 1 and 3, 5 and 7). These results indicate that while a majority of P-TEFb (CDK9:CycT1) is regulated via 7SKsnRNP equilibrium, a majority of P-TEFb (CDK9:CycT2) is free from 7SKsnRNP, and 7SKsnRNP-associated P-TEFb (CDK9:CycT2) responds to JQ1 to a lesser extent than P-TEFb (CDK9:CycT1).

### Transcriptome analysis reveals that expression of inducible, but not steady-state, genes are affected in CycT1-KO cells

Next, to determine whether basal- and P-TEFb-induced transcription is altered in CycT1-KO cells, we compared transcriptomes between WT and CycT1 KO cells with or without JQ1 stimulation. Cells were left unstimulated or stimulated with JQ1 to induce P-TEFb for 12hr, and whole cellular RNAs were isolated and subjected to RNA-seq analysis (**Fig. 4A**). A rather small difference between unstimulated WT and CycT1-KO cells was observed, and changes in the levels of most genes were within 2-fold (**Fig.4B**). This indicates that lack of CycT1 has a minor impact on the steady-state transcription. Next, JQ1-dependent transcription (ratio of JQ1+/JQ1-) of each gene in WT and CycT1(-) cells were calculated and plotted (**Fig. 4C**, ALL GENES), which indicates that changes in the expression levels of many genes were also within 2 folds between WT and CycT1-KO cells. We further divided the genes into three categories based on JQ1 responsiveness in WT cells: (1) genes up-regulated by JQ1 (log2FC>2), (2) genes unaffected by JQ1 (2>log2FC>-2), and (3) genes down-regulated by JQ1 (log2FC<-2). Expression levels of genes in each category were compared between WT and CycT1 (-) cells. As shown in **Fig.4C**, the levels of JQ1-up-regulated genes were significantly lower in CycT1-KO cells, compared with WT cells (**Fig.4C**, “UPREGULATED”). A similar tendency was observed with JQ1-down-regulated genes (**Fig.4C**, “DOWNREGULATED”). The mechanism of JQ1-dependent transcriptional suppression is still largely unknown, and it is possible that the suppression could be a result of P-TEFb-dependent upregulation of negative factors such as HEXIM1 [20, 21] . There was no difference in UNCHANGED genes between WT and CycT1-KO cells (**Fig.4C**, “UNCHANGED”). The scatter blot between changes in gene expression by JQ1 between WT and CycT1-KO cells indicates that most JQ1-dependent genes are affected in CycT1-KO cells (**Fig.4D**). These results are consistent with the results of reporter gene assays (**Fig.3A**). Next, among the JQ1-up-regulated genes, top thirteen genes that responded to JQ1 to the highest extent in WT cells were depicted and compared between WT and CycT1-KO cells (**Fig. 5A**). The name and function of each gene product is summarized in **Fig.5B**. It is to be noted that HEXIM1 was among these genes, which is consistent with previous studies that HEXIM1 is one of the immediately early genes activated by P-TEFb stimulation [16, 21]. As expected, most genes were not upregulated by JQ1 in CycT1-KO cells (**Fig. 5A**, gray bars compared with black bars). These results indicated that CycT1 is required for expression of the genes that are highly inducible by JQ1.

**Figure 3.**
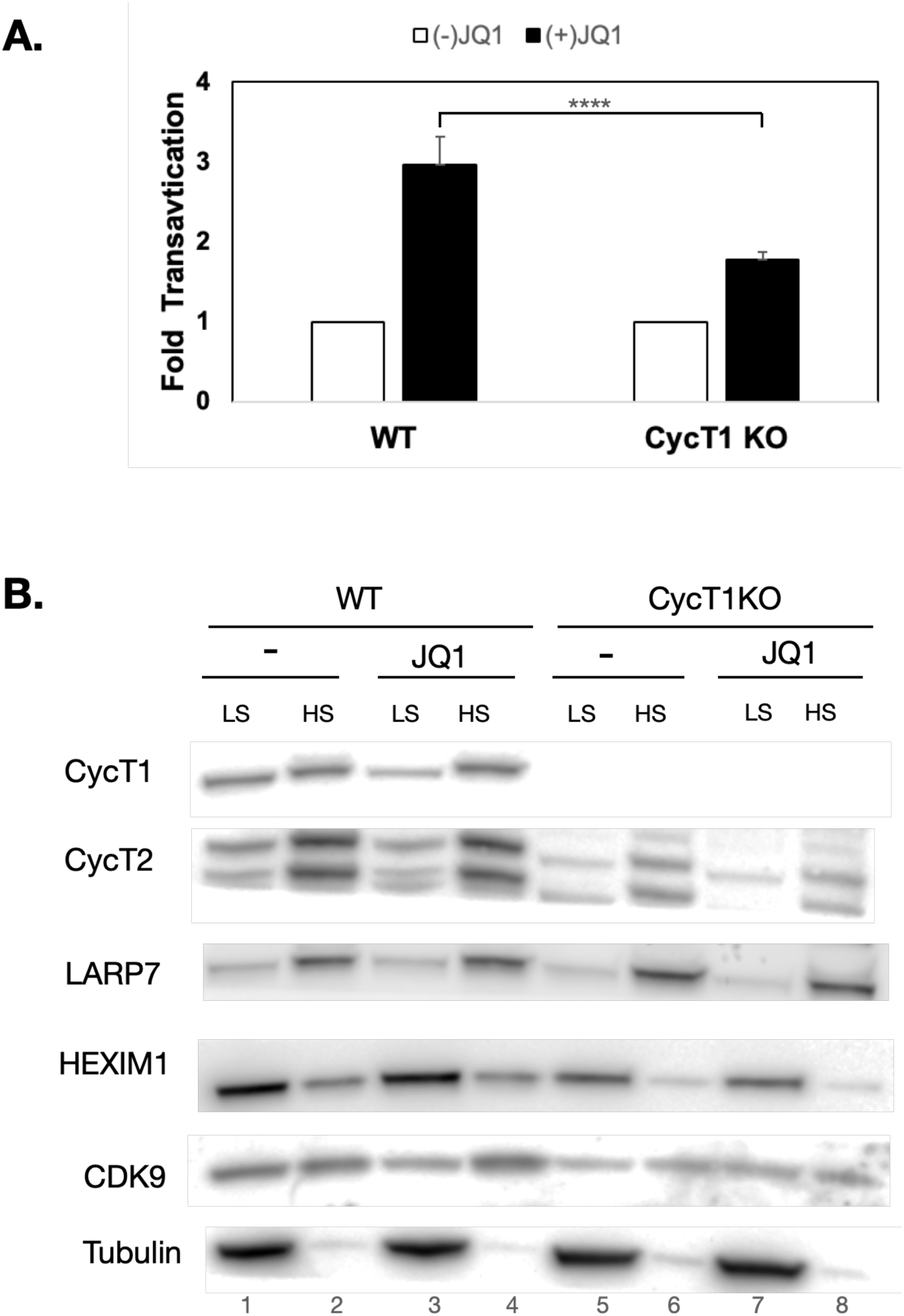
JQ1-induction of P-TEFb is impaired in CycT1-KO cells. **A.** 293T or CycT1-KO cells were transfected with a luciferase reporter plasmid that containing the promoter region of the HEXIM1 gene (HexP-Luc). 24 hours after transfection, cells were treated with DMSO or JQ1 for additional 24 hours. Cells were then harvested and subjected to luciferase assays. Results are presented as values relative to that obtained with 293T cells treated with DMSO. **B.** 293T or CycT1-KO cells were treated with JQ1 for 2 hours. Cells were harvested and lysed in the low salt (LS) lysis buffer, and the supernatants were set aside. The cell pellets were then lysed in the high salt (HS) buffer. Both LS and HS fractions were separated by SDS-PAGE followed by Western blotting with the antibodies indicated at the left of each panel.

**Figure 4.**
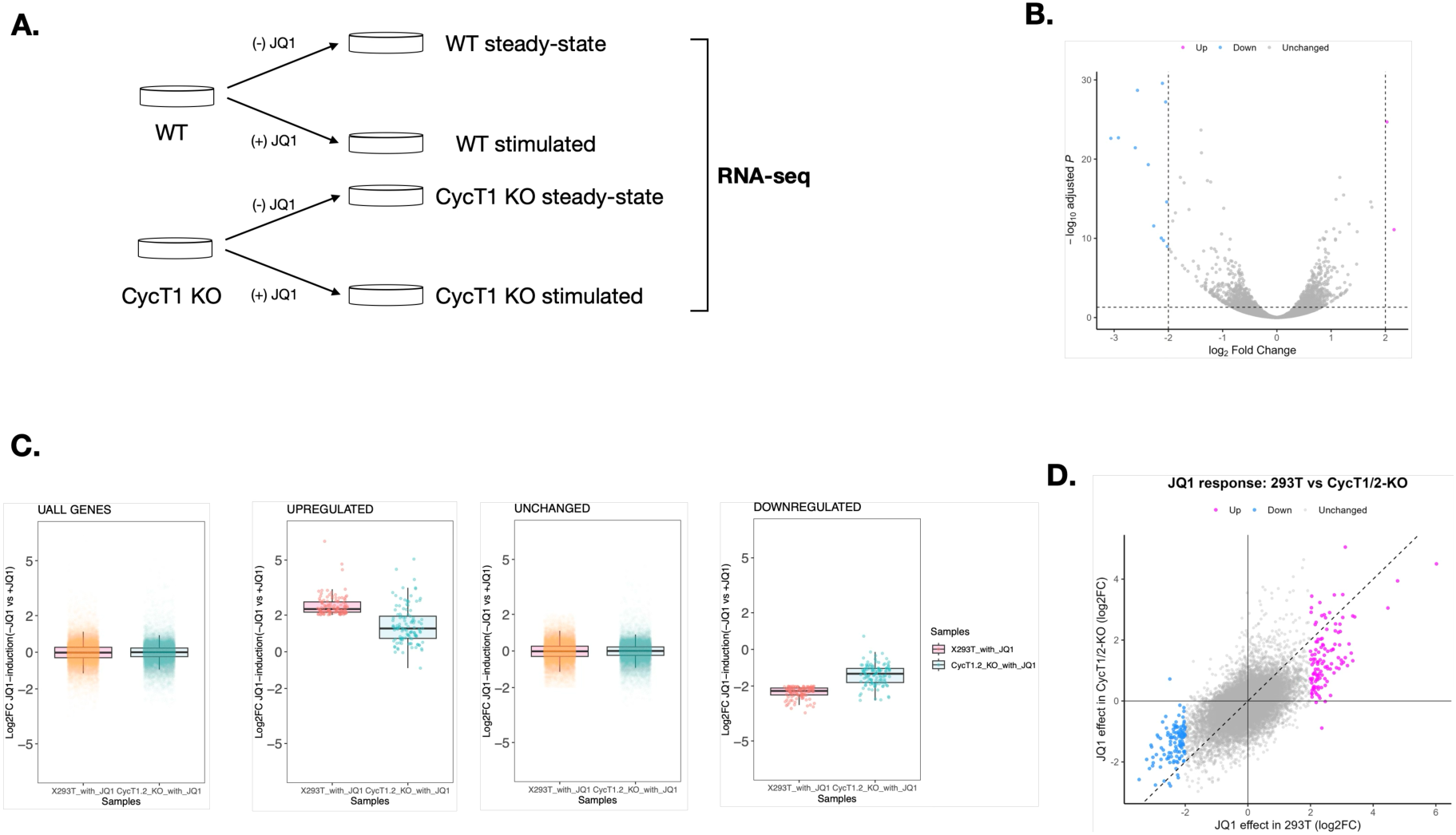
Global gene expression analysis of CycT1-KO cells. **A.** A schematic presentation of the RNA-seq analysis. 293T and CycT1-KO cells were untreated or treated with JQ1 for 12 hours. Total RNA was isolated and subjected to RNA-Seq. **B.** The volcano plot shows differential gene expression for JQ1 treatment in 293T cells, JQ1 treatment in CycT1/2-KO cells, and CycT1/2-KO relative to 293T cells. The x-axis represents log2 fold change (log2FC), and the y-axis represents −log10 adjusted *P* value. Genes with an adjusted *P* value < 0.05 and log2FC ≥ 2 or ≤ −2 were classified as upregulated (magenta) or downregulated (blue), respectively; all other genes are shown in gray. **C.** All genes were categorized in three groups based on the difference in expression levels between unstimulated and JQ1-stimulated 293T cells: genes upregulated by JQ1, genes unchanged (-2<log2,2), and genes downregulated by JQ1. In each group, differences in mRNA expression levels of each gene in between JQ1-stimulated 293T cells and JQ1-stumulated CycT1-KO cells are plotted. **D.** The scatter plot compares the effect of JQ1 in 293T cells (x-axis) with that in CycT1/2-KO cells (y-axis). Genes are classified and colored according to their JQ1 response in 293T cells using the same criteria. The dashed diagonal line indicates y = x.

**Figure 5.**
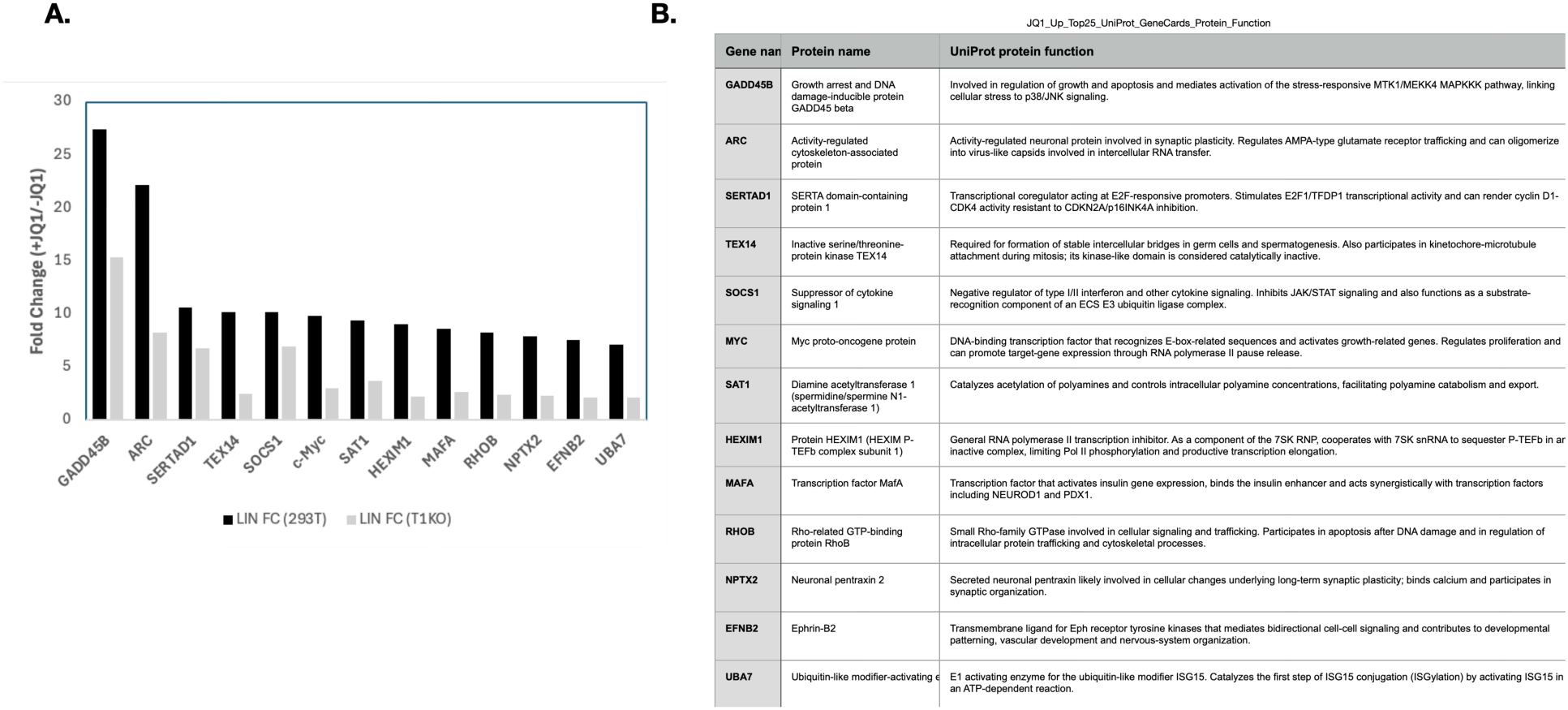
A. Top 13 JQ1-upregulated genes in 293T were depicted and compared expression levels in between 293T and CycT1-KO cells. **B.** Names and functions of the selected genes are summarized in a table. Functions of the selected genes are based on the description in Uniplot database.

### JQ1-inducible genes affected in CycT1-KO cells are upregulated in Pancreatic Adenocarsioma (PAAD)

These thirteen genes were further analyzed using publicly available database of gene expression in various types of cancer (https://oncodb.org), which provides integrated multi-omic data for approximately 10,000 patients across 33 cancer types. Each gene of interest was searched in all 33 cancer types and examined whether the gene is upregulated or downregulated in each cancer type with statistical significance.

Interestingly, 11 out of 13 CycT1-/JQ1-dependent genes are upregulated in Pancreatic Adenocarcinoma (PAAD) (**Fig. 6**). In particular, the expression of HEXIM1, SERTAD1, EFNB2, RHOB, SAT1, and UBA1 is upregulated with high significances (P<1.0e-30, **Fig.6B, G, I, J, K, M**). All other P-TEFb/7SKsnRNP components except for CycT1 were also found upregulated in PAAD (**Fig.7**) although it is unclear whether P-TEFb activity is augmented in PAAD. These results suggest that genes highly inducible by JQ1 in CycT1-dependent manner are involved in certain types of cancers.

**Figure 6.**
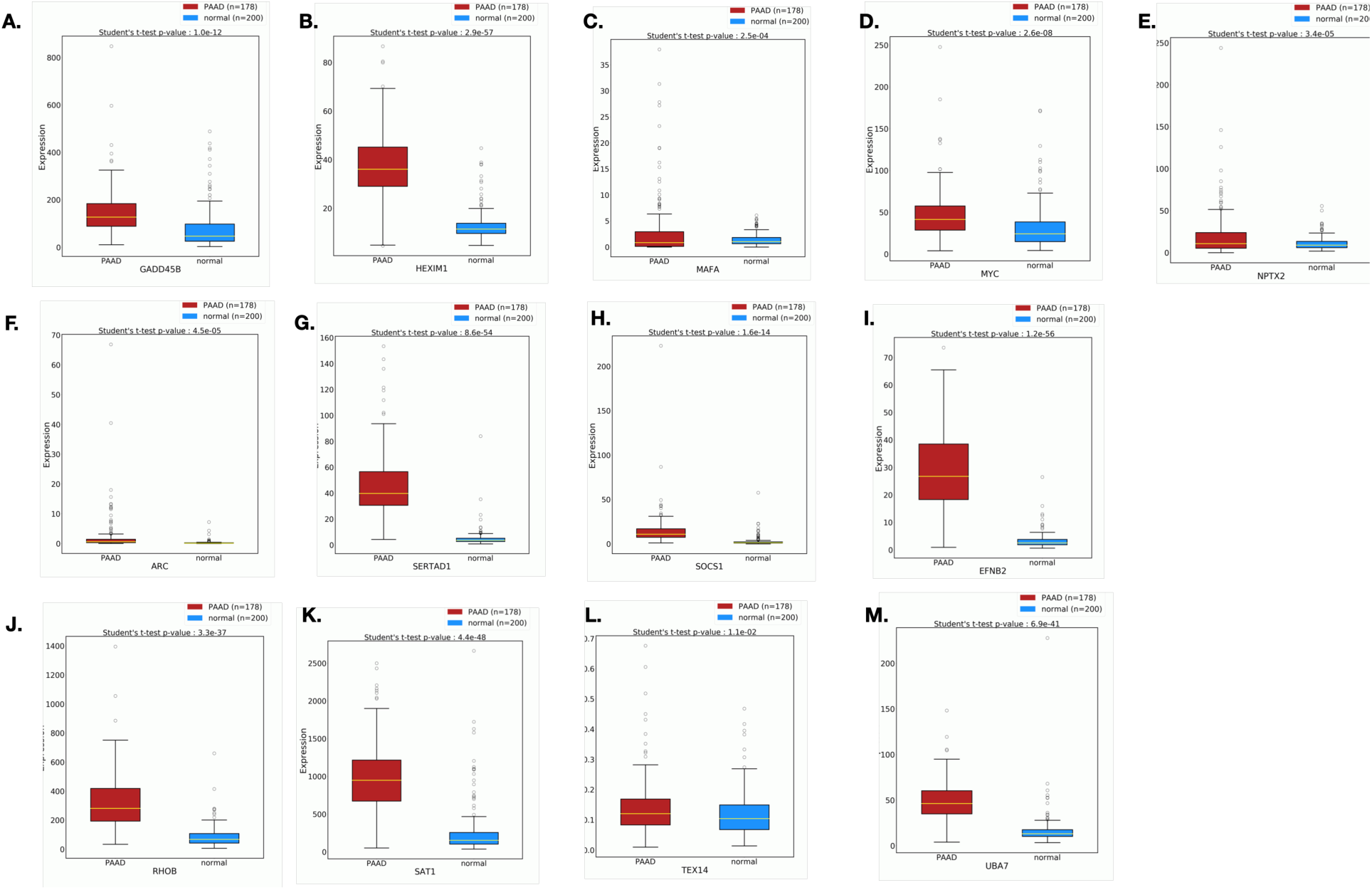
JQ1-upregulated/CycT1-dependent genes were also upregulated in pancreatic adenocarcinoma (PAAD). The 13 genes depicted in Fig.4D were searched in OncoDB database to obtain gene expression data in different types of cancer. 11 out of 13 genes were found upregulated in 178 specimens of PAAD. Gene expression of PAAD (red) and normal (blue) specimens were presented by box plot graphs.

**Figure 7.**
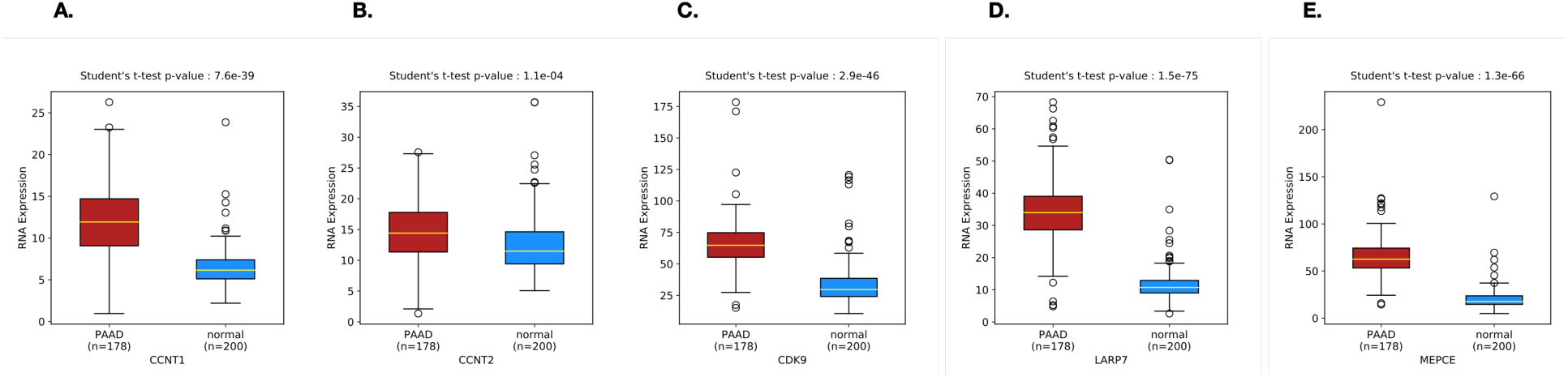
Gene expression analysis of P-TEFb and 7SKsnRNP components in PAAD. Using the OncoDB database, the expression of CCNT1 (**A**), CCNT2 (**B**), CDK9 (**C**), LARP7 (**D**) and MEPCE (**E**), were compared between PAAD (red) and normal (blue) specimens. Expression levels of 7SKsnRNA (long non-coding RNA) were unable at OncoDB.

## Discussion

To analyze the role of the CycT1 subunit of P-TEFb on cellular transcription, we stablished HEK293T cells lacking CycT1 proteins (CycT1-KO cells). No apparent growth defects were observed with CycT1-KO cells. Although level of CDK9 was decreased and no compensatory over-expression of CycT2 occurred, a majority of P-TEFb (CDK9:CycT2) are not incorporated in 7SKsnRNP complexes. NFκB-dependent transcription, but not AP1-dependent transcription, induced by PMA was impaired in CycT1-KO cells although the function of NFκB (P65) *per se* was not affected. Moreover, transactivation by JQ1 was impaired in CycT1-KO cells. JQ1 released P-TEFb (CDK9:CycT1) while majority of P-TEFb (CDK9:CycT2) was not affected by JQ1.

Transcriptome analysis indicates that the lack of CycT1 had a minor effect on the steady-state transcription. However, stimulation of JQ1-dependent genes was severely impaired in CycT1-KO cells. Finally, data mining of transcriptome databases revealed that 11 out of 13 highly inducible JQ1-dependent genes were also upregulated in Pancreatic Adenocarcinoma. From these results, we conclude that although many genes can be regulated by both P-TEFb (CDK9:CycT1) and P-TEFb (CDK9:CycT2), CycT1 plays a critical role in regulating highly inducible genes, which are also aberrantly regulated in a particular type of cancer.

To our knowledge, this study is the first example of genetically eliminating CycT1 subunit of P-TEFb to obtain a comprehensive view of CycT1-dependent transcription. Although CycT1 is a major cyclin to form P-TEFb complexes in most cell types, complete elimination of CycT1 did not affect cells’ viability, suggesting CycT2s can functionally compensate the loss of CycT1 for the normal cell growth. While the level of CycT2 did not increase in CycT1-KO cells, majority of the P-TEFb (CDK9:CycT2) complexes are free from 7SKsnRNP and possibly engaged on RNAPII on genes required for the cellular homeostatis. Our results suggest that P-TEFb (CDK9:CycT1) is particularly important for inducible transcription, for which P-TEFb (CDK9:CycT2) cannot fully compensate the loss of CycT1. Although transcription induced by PMA or JQ1 was impaired in CycT1-KO cells, the precise mechanisms are still yet to be determined. For the JQ1 stimulation, while P-TEFb (CDK9:CycT1) complexes are translocated from the LS fraction (cytoplasmic or loosely associated with the nucleus) to the HS fraction (tightly associated to the nucleus) after JQ1 stimulation, the distribution of P-TEFb (CDK9:CycT2) between these two fractions are unaffected by JQ1. Further studies will be required for elucidating precise molecular mechanisms.

Although no human diseases are associated with genetic mutations in the CCNT1 gene, levels of CycT1 proteins are often correlated with specific types of cells. For example, in resting and unresponsive T cells, levels of CycT1 proteins are vanishingly low, and as a consequence, amounts of the functional P-TEFb complexes are kept at very low levels in these cells [22, 23]. Our CycT1-KO cells partially mimic these conditions. Our previous studies demonstrated that CycT1 proteins are dephosphorylated by protein phosphatase (PP)1 in resting CD4+ T cells, which dissociates CycT1 from CDK9.

CDK9-unbound CycT1 proteins are rapidly degraded SIAH1/2 E3 ligase-dependent proteasomal pathways. In replicating cells, on the other hand, CycT1 proteins are phosphorylated at positions 143 (T143) and 149 (T149) by PKC, which potentiates the interaction between CycT1 and CDK9 and stabilize P-TEFb complexes [10, 12]. Lack of P-TEFb (CDK9:CycT1) complexes in quiescent CD4+ T cells is a main cause of the latent infection of HIV-1 [24]. Since our data suggest P-TEFb (CDK9:CycT1) is required for signal-induced transcription, the lack of functional P-TEFb (CDK9:CycT1) would be a main cause of impaired cytokine production by T cell receptor signaling in anergic and/or exhausted T cells. CycT1-KO cells would therefore be useful tools to study the mechanism of signal-induced transcription activation and develop methods to restore T cell responses.

While functions of CycT1 have been extensively studied, roles of CycT2 are currently underexplored. Nevertheless, several studies have suggested CycT2 plays critical roles during various developmental stages. CycT2 is essential for mouse embryogenesis since deletion of the CCNT2 gene is embryonic lethal. Other studies indicate that CycT2 is required for muscle differentiation via interacting with the MyoD transcription factor, early spermatogenesis, or developmental programming of adipose tissues [7, 25–27].

Those studies are mainly based on differences in mRNA expression patterns of CCNT1 and CCNT2 genes, and it is still largely unclear whether P-TEFb (CDK9:CycT1) and P-TEFb (CDK9:CycT2) activate different target genes via distinct pathways. Our results provide important clues to determine the mechanisms of redundant and non-redundant gene expression by different P-TEFb complexes.

Pancreatic Adenocarcinoma (PAAD) is a most common and aggressive type of pancreatic cancer, reaching 95% of the cases of pancreatic cancer. Also, most (>90%) PAAD are driven by mutations in the KRAS genes. Recently a pan-RAS inhibitor, daraxonrasib, made a breakthrough by drastically increasing life expectancies of patients with PAAD [28, 29]. At this point, it is yet to be determined whether P-TEFb activity was aberrantly upregulated in PAAD, although mRNA expression of CycT1, CDK9, LARP7, and MEPCE, but not CycT2, are increased in PAAD. Also, whether the 11 highly P-TEFb-dependent genes play critical roles in PAAD is unclear. Interestingly, one of the commonly used compounds for the treatment of pancreatic adenocarcinoma is 5-FU [30]. We have previously demonstrated that 5-FU also induces the P-TEFb release from 7SKsnRNP. Many P-TEFb inducers increase HEXIM1 expression, which in turn causes global transcriptional repression via re-incorporation of P-TEFb into 7SKsnRNP complexes. 5-FU might also cause global transcription repression after the initial upregulation of P-TEFb activities. It is therefore of interests whether blocking P-TEFb enhances the effect of daraxonrasib.

## Materials and Methods

### Cell lines, antibodies and plasmids

Human Embryonic Kidney (HEK) 293T cells were cultured in Dulbecco’s modified Eagle’s medium (DMEM) (Corning) with 10% fetal bovine serum (FBS) (Sigma Aldrich). Antibodies used in this study for co-immunoprecipitations and western blotting were: α-HEXIM1 (Hex)(SBCT, )(Proteintech, 15676-1-AP), α-CDK9 (SCBT,F6)(Proteintech, 11705-1-AP), α-CycT1 (SBCT, sc-2713483)(Proteintech, ), α-LARP7 (Proteintech), α-MEPCE (Proteintech), α-tubulin (Proteintech 11224-1-AP), and α-Flag (Sigma Aldrich, F-3165). HexP-, UAS- and HIV-LTR luciferase reporter plasmids were described elsewhere (ref). NFκB- and AP1-Luciferase reporter plasmids were kind gifts from Dr. Yang Li (UCSF).

### Co-immunoprecipitation (Co-IP)

293T cells were lysed with chilled RIPA buffer (50 mM Tris-HCl, pH 8.0, 5 mM EDTA, 0.1% SDS, 1.0% Nonidet P-40, 0.5% sodium deoxycholate, 150 mM NaCl) containing protease inhibitors (Rioche). The supernatant was precleared with protein G-Sepharose beads for 2 h. The precleared supernatant was incubated with indicated primary antibodies or control IgG pre-coupled with protein G-Sepharose beads overnight. Beads were washed 5 times with the RIPA buffer. The co-IP samples and input (5% of whole cell lysates) were boiled in 2× Laemmli sample buffer (Bio-Rad) supplemented with 2-mercaptoethanol (Bio-Rad) and subjected to western blotting (WB) as described previously [19].

### Sequential salt extraction

Differential salt extraction was carried out to determine fractions of free P-TEFb or 7SK snRNP according to Biglione et al., with some modifications [15, 32, 33]. Cells were collected and washed twice with cold PBS. Cells were lysed in 200 µl low salt buffer (10 mM KCl, LS, 10 mM MgCl_2_, 10 mM HEPES-KOH pH 7.5, 1 mM EDTA, 1mM DTT, 0.5% NP40, proteinase inhibitor cocktail) and incubated on ice for 10 min. Lysates were then centrifuged at 5000 X g for 5 min, and supernatants were collected and designated as 7SK snRNP fractions. Pellets were washed once with 200 µl LS buffer and resuspended in 70 µl high salt buffer (450 mM NaCl, HS, 1.5 mM MgCl_2_, 20 mM HEPES pH7.5, 0.5 mM EDTA, 1 mM DTT, 0.5% NP40, proteinase inhibitor cocktail). Lysates were then centrifuged at 10000 X g for 5 min, supernatants were collected and diluted x3 with LS buffer to adjust the salt concentration.

### CRISPR-Cas9-based genetic inactivation

CycT1 KO cells were created using the CRISPR/Cas9 system. All-in-one Cas9/sgRNA plasmid pSpCas9 BB-2A-Puro encoding sgRNA specific for CycT1 (sgRNA sequence: AAGCAGATTGGCCGCCTGC) (GenScript) was transfected into 293T cells using Lipofectamine 2000 (Invitrogen). After 48 hrs, untransfected cells were selected against by puromycin treatment for another 72 hrs. After puromycin selection, cells were cultured in normal media, and further cloned by limiting dilution. CycT1 KO clones were selected by analyzing CycT1 protein expression by WB using at least two different anti-CycT1 Abs (SCBT SC10750 and SC271348). Genomic CycT1 sequences were also analyzed to confirm that both alleles contained mutations that cause truncation of CycT1 protein at the N-terminus.

### Luciferase assays

Luciferase reporter plasmids (20-50ng) were transfected using Lipofectamine 2000 (Invitrogen) with or without mammalian expression plasmids encoding indicated proteins (20-100ng) in 293T or CycT1KO cells (triplicate with 1E+5 cells in 96 well plates). After 24hrs, cells were untreated or treated with DMSO, the cells were washed using PBS, lysed with Passive Lysis Buffer (Promega), and analyzed for Luciferase expression using D-Luciferin (BD Monolight) on an EG&G Berthold LB 96V microplate luminometer.

### mRNA-seq

1E+06 293T or CycT1-KO cells were treated with DMAO or JQ1 (5μM) for 12 hours. Cells were washed once with PBS, and total cellular RNA was purified RNeasy Mini kit (Qiagen), followed by DNase I treatment (Qiagen). 500 ng RNA was submitted for RNA-seq analysis at UCSF’s Functional Genomics Core Facility. Each condition was tested in triplicate experiments.

### Data mining

CycT1- and JQ1-dependent genes were further analyzed at OncoDB database (https://oncodb.org), offering integrated multi-omic data for approximately 10,000 patients across 33 cancer types. Each gene of interest was searched in all of 33 cancer types and examined whether the gene is upregulated or downregulated in each cancer type with statistical significance.

### Statistical analysis

For p24 ELISA, luciferase enzymatic assays, GFP assays, microscopic analyses and RT-qPCR measurements, 3 independent experiments were performed in duplicate.

Thus, bars represent SEM, n=3. A Student’s *t*-test was preformed to measure the significance of the data (* P<0.05, ** P<0.01, *** P<0.001).

